# Enhanced detection of nucleotide repeat mRNA with hybridization chain reaction

**DOI:** 10.1101/2021.01.06.425640

**Authors:** M. Rebecca Glineburg, Yuan Zhang, Elizabeth Tank, Sami Barmada, Peter K Todd

## Abstract

RNAs derived from expanded nucleotide repeats form detectable foci in patient cells and these foci are thought to contribute to disease pathogenesis. The most widely used method for detecting RNA foci is fluorescence in situ hybridization (FISH). However, FISH is prone to low sensitivity and photo-bleaching that can complicate data interpretation. Here we applied hybridization chain reaction (HCR) as an alternative approach to repeat RNA foci detection of GC-rich repeats in two neurodegenerative disorders: GGGGCC (G_4_C_2_) hexanucleotide repeat expansions in *C9orf72* that cause amyotrophic lateral sclerosis and frontotemporal dementia (C9 ALS/FTD) and CGG repeat expansions in *FMR1* that cause Fragile X-associated tremor/ataxia syndrome. We found that HCR of both G_4_C_2_ and CGG repeats has comparable specificity to traditional FISH, but is >40x more sensitive and shows repeat-length dependence in its intensity. HCR is better than FISH at detecting both nuclear and cytoplasmic foci in human C9 ALS/FTD fibroblasts, patient iPSC derived neurons, and patient brain samples. We used HCR to determine the impact of integrated stress response (ISR) activation on RNA foci number and distribution. G_4_C_2_ repeat RNA did not readily co-localize with the stress granule marker G3BP1, but ISR induction increased both the number of detectible nuclear RNA foci and the nuclear/cytoplasmic foci ratio in patient fibroblasts and patient derived neurons. Taken together, these data suggest that HCR can be a useful tool for detecting repeat expansion mRNA in C9 ALS/FTD and other repeat expansion disorders.

## INTRODUCTION

GC-rich repeat expansions are the genetic cause of over 50 neurodevelopmental, neurodegenerative, and neuromuscular diseases. Repeat expansions elicit disease through multiple mechanisms (*reviewed in* Orr and Zoghbi 2007; Rodriguez and Todd 2019). Repeats as DNA can inhibit transcription of repeat-containing genes and promote the DNA damage response through R-loop formation (Tuduri et al. 2009; Reddy et al. 2011; Reddy et al. 2014; Groh et al. 2014; Haeusler et al. 2014). Repeats as RNA can functionally sequester both canonical and non-canonical RNA binding proteins and prevent them from performing their normal functions (Miller et al. 2000; Mankodi et al. 2000; Jin et al. 2007; Cooper-Knock et al. 2014; Conlon et al. 2016; Lee et al. 2013; reviewed in Rodriguez and Todd 2019; Swinnen, Robberecht, and Van Den Bosch 2020; Johnson and Cooper 2021). Repeats can also be translated into toxic proteins, either via canonical translation (as occurs for a number of polyglutamine expansions) or via repeat associated non-AUG (RAN) initiated translation (Zu et al. 2011; Todd et al. 2013; Green et al. 2017; Kearse et al. 2016; Paulson et al. 1997; Trottier et al. 1995; La Spada et al. 1991).

One of the key pathological hallmarks in repeat expansion diseases is the presence of repeat-containing RNA foci. These foci are thought to represent repeat RNAs co-associated with specific RNA binding proteins, although they may also arise from intra- and inter- molecular RNA-RNA interactions via RNA gelation (Jain and Vale 2017; Fay, Anderson, and Ivanov 2017; Mankodi et al. 2001; Fardaei et al. 2002; Tassone, Iwahashi, and Hagerman 2004; Iwahashi et al. 2006; Jin et al. 2007; Sofola et al. 2007; Sellier et al. 2010; Sellier et al. 2013; Bajc Cesnik et al. 2019). The exact behavior, biophysical properties, and associated protein and RNA factors of these RNA condensates varies across different repeat expansions *(reviewed in* Nussbacher et al. 2019; Echeverria and Cooper 2012; Johnson and Cooper 2021). Traditionally, these foci are visualized by fluorescent in situ hybridization (FISH) (Figure 1A) (Urbanek and Krzyzosiak 2016; Lee et al. 2013; Urbanek, Michalak, and Krzyzosiak 2017; Hoem et al. 2011; Verma et al. 2019; Jin et al. 2007). Because of the repetitive nature of the repeat, single probes to the repeat region can “tile” along the mRNA and enhance signal detection, early studies were able to successfully identify these foci. However, FISH in human tissues has remained hindered by the low abundance of these repeat containing RNAs, and high background due to the human transcriptome containing numerous GC rich repeats (Sulovari et al. 2019; Saxonov, Berg, and Brutlag 2006; Yamashita et al. 2005; Kozlowski, de Mezer, and Krzyzosiak 2010; DeJesus-Hernandez et al. 2011; Belzil et al. 2013; Urbanek and Krzyzosiak 2016; Didiot et al. 2018).

**Figure 1.**
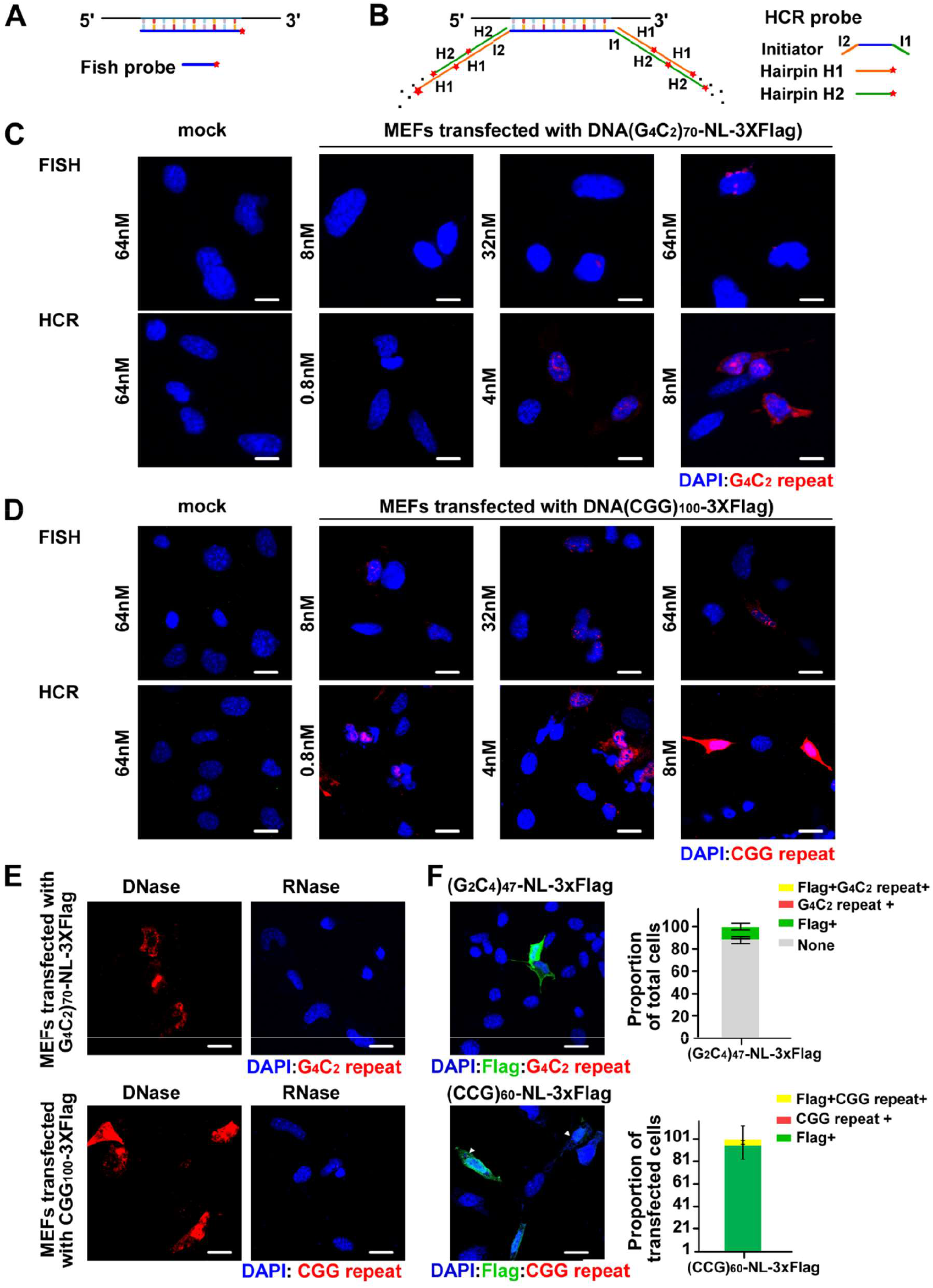
Hybridization Chain Reaction mediated detection of GC rich repeats. **A)** FISH probe with a 5’ conjugated fluorophore. Signal strength is dependent on number of RNA molecules present. **B)** In HCR, three probes are used: the initiator binds directly to the RNA of interest and has both 3’ and 5’ extensions complementary to two 5’ fluorophore conjugated hairpin probes (H1 and H2). Upon binding, H1 and H2 unfold to reveal new binding sites for the other hairpin probe. In this way, signal from one RNA molecule is amplified > 100 fold, dramatically enhancing detection. **C-D)** MEFs transfected with (G_4_C_2_)_70_-NL-3xFlag (C) or CGG_100_-3xFlag (D) vectors and probed with indicated concentrations of either FISH or HCR probe. **E)** MEFs transfected with (G_4_C_2_)_70_-NL-3xFlag expressing vector (top) or CGG_100_-3xFlag vector (bottom) and treated with DNase (left) or RNase (right) prior to HCR. **F)** MEFs transfected with antisense (G_2_C_4_)_47_ (top) or (CCG)_60_ (bottom) expressing plasmids followed by ICC stained with Flag and HCR sense strand probes (C_4_G_2_)_6_ or (CCG)_8_. Top: N=434; Bottom: N=38. Error bars indicate 95% confidence intervals (CI). Scale bar=20 μ m in C, D, E; 50 μ m in F.

Recently, a more sensitive method, hybridization chain reaction (HCR) was developed that uses an initiator probe recognizing the RNA of interest and a pair of hairpin probes conjugated with fluorophores to amplify the initiator signal (Figure 1B) (Choi et al. 2018; Choi, Beck, and Pierce 2014a, 2014b; Choi et al. 2011). This approach significantly amplifies the signal of an individual molecule over traditional FISH and makes it easier to detect low abundant RNAs, while also reducing background signal. Here, we utilized HCR to detect RNA foci associated with GGGGCC repeat expansions in C9orf72 that are the most common genetic cause of ALS and FTD and CGG repeats associated with Fragile X disorders such as Fragile X-associated tremor/ataxia syndrome (FXTAS) (Hagerman et al. 2001; Renton et al. 2011; DeJesus-Hernandez et al. 2011). HCR provided significantly higher sensitivity for detection of repeats and allowed accurate tracking of endogenous GGGGCC repeat RNAs in patient cells in response to cellular stress activation. Taken together, these data suggest that HCR can be a valuable addition to analysis and imaging pipelines in repeat expansion disorders.

## Results

### HCR is more sensitive than FISH for GC rich repeats

We first compared the sensitivity and specificity of traditional FISH vs HCR probes. We designed fluorophore labeled locked nucleic acid (LNA) (C_4_G_2_)_6_ and (CCG)_8_ FISH probes and (C_4_G_2_)_6_ and (CCG)_10_ HCR initiator probes to hybridize to corresponding G_4_C_2_ or CGG repeats. Alexa fluorophore labeled amplifier hairpin probes B1H1 and B1H2 were then used to amplify the HCR probe signal (Figure 1 A-B, Supplemental table 2) (Choi, Beck, and Pierce 2014b; Choi et al. 2011). We conducted a side-by-side comparison in mouse embryonic fibroblasts (MEFs) expressing (G_4_C_2_)_70_-NL-3xFlag or (CGG)_100_-3xFlag reporters (Green et al. 2017; Kearse et al. 2016). We found more repeat positive cells by HCR than FISH when using the same probe concentration (8nM) (Figure 1C-D). When we further increased each FISH probe concentration to 64nM, we only saw a modest increase in number of repeat positive cells that remained significantly lower than that seen using HCR. In contrast, when we decreased each HCR probe concentration to 4nM and 0.8nM, we still observed enhanced signal compared to FISH, for both G_4_C_2_ and CGG repeats (Figure 1C-D). Furthermore, both nuclear and cytoplasmic signal was readily detected using HCR, while FISH primarily detected only the stronger nuclear signal. No signal was detected by either method in MEFs not expressing G_4_C_2_ or CGG repeats (Figure 1C-D). Together, these results show that HCR is more sensitive than FISH for detecting exogenous GC rich repeats.

To confirm that the observed signal from HCR was due to probe hybridization to RNA and not DNA, we treated the (G_4_C_2_)_70_ and (CGG)_100_ transfected MEFs with DNase or RNase prior to HCR. While DNase robustly eliminated DAPI signal, it had no effect on the GC-rich repeat signal. Conversely, almost all probe signal went away when cells were treated with RNase (Figure 1E). To further validate that our HCR probes were specifically recognizing their target RNA, we transfected cells with antisense CCCCGG (ATG-(C_4_G_2_)_47_-NL-3xFlag) or CCG ((CCG)_60_-NL-3xFlag) reporters. We did not detect any repeat signal in Flag positive cells (Figure 1F). Together this supports that our HCR probes specifically hybridize to G_4_C_2_ and CGG repeat containing RNA, with high specificity and sensitivity.

We next determined whether HCR could distinguish between different G_4_C_2_ repeat sizes (3, 35, or 70 G_4_C_2_ repeats). Despite similar transfection efficiencies among all three conditions (as monitored by GFP co-transfection, Supplemental Figure 1A-C), we observed no signal in cells expressing (G_4_C_2_)_3_-NL-3xFlag. The number of HCR positive cells was comparable for both (G_4_C_2_)_35_-NL-3xFlag and (G_4_C_2_)_70_-NL-3xFlag transfection (Figure 2A-B). However, there was a significant correlation between HCR signal intensity and repeat length, with (G_4_C_2_)_70_-NL-3xFlag transfected cells having higher HCR signal intensity (Figure 2C).

**Figure 2.**
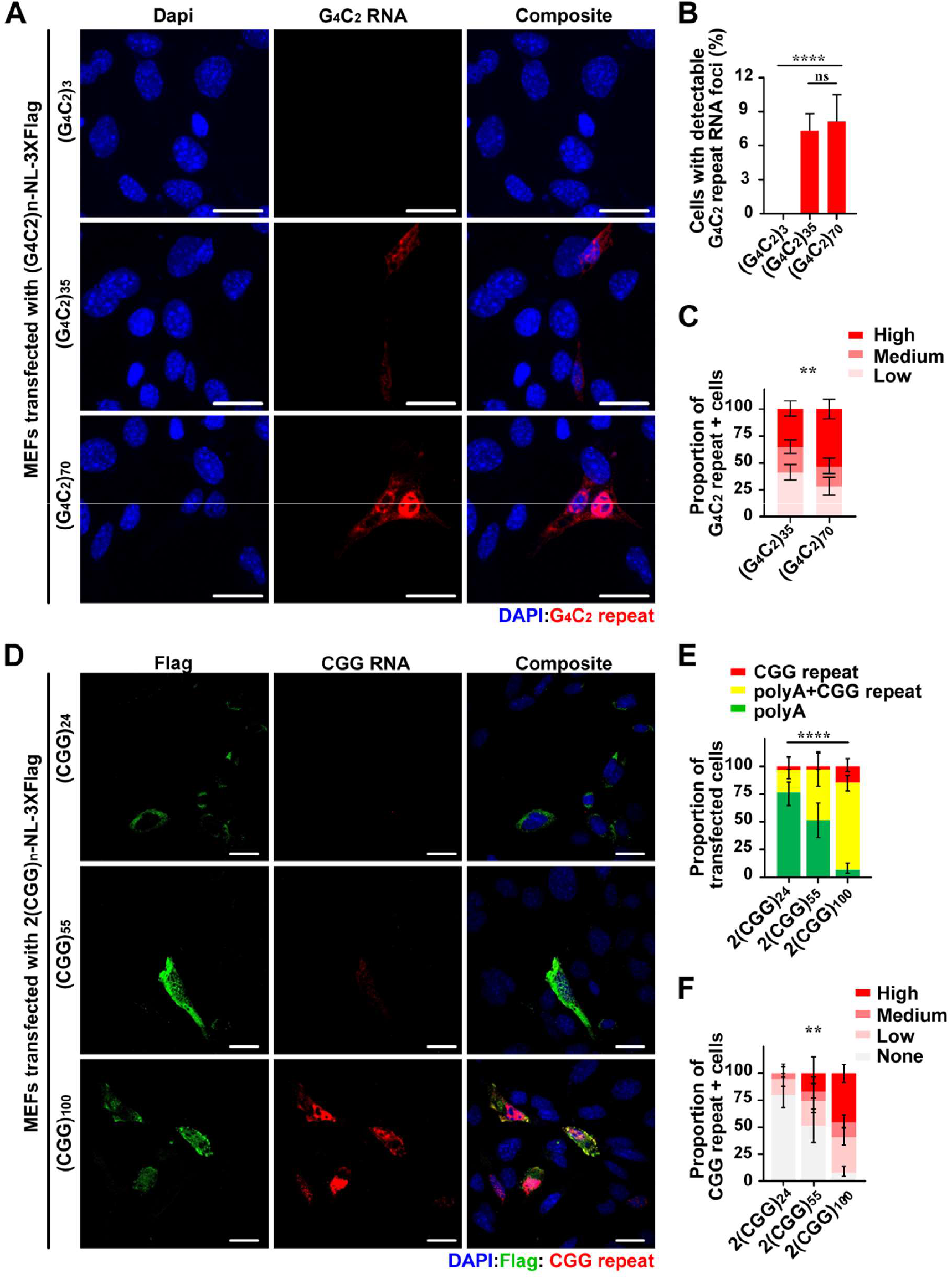
HCR probe signal intensity is repeat length dependent for G_4_C_2_ and CGG repeats. **A)** MEFs transfected with G_4_C_2_ repeat constructs with increasing repeat sizes. **B)** Quantification of cells with detectable G_4_C_2_ repeat RNA foci in total cells. (G_4_C_2_)_3_: N=313; (G_4_C_2_)_35_ N=575; (G_4_C_2_)_70_ N=514. **C)** HCR signal intensity in MEFs transfected with (G_4_C_2_)_n_-NL-3xF reporters with indicated repeat length. (G_4_C_2_)_35_: N=175; (G_4_C_2_)_70_: N=110. **D)** MEFs transfected with CGG repeat constructs with increasing repeat sizes. **E)** Quantification of Flag+ and CGG RNA+ cells expressed as proportion of total transfected cells. **F)** HCR signal intensity in MEFs transfected with 2(CGG)n-NL-3xF reporters with indicated repeat length. E-F) N ≥35/condition. Error bar indicates 95% CI. Statistics: one-way ANOVA for B, Chi-square test for C, E and F. \*\**p*<0.01, \*\*\*\**p*<0.0001, ns: not significant. Scale bar=50 μ m in A and D.

We next performed similar experiments in cells transfected with CGGn-NL-3xFlag reporters with repeats of various length (24, 55, and 100 CGG repeats). We observed a strong positive correlation between CGG repeat length and the number of HCR positive cells (Figure 2D-E). We also observed a significant increase in HCR signal intensity within cells expressing longer CGG repeats (Figure 2D & F). CGG repeat expansions within the *FMR1* 5’UTR have previously been shown to enhance transcription, and we and others have observed a positive correlation of repeat length and RNA abundance in this setting (Chen et al. 2003; Khateb et al. 2007 and unpublished data). Thus, this repeat-length increase in HCR signal could result from enhanced mRNA production as well as more CGG repeat binding sites in the longer repeat reporters. As both repeat size and expression are tightly linked to disease, we believe HCR can be used to qualitatively assess CGG repeat RNA burden in model systems.

### HCR is more sensitive in detecting endogenous repeat signal than FISH

We next determined if HCR was effective at detecting endogenous GC-rich repeat RNA. We first compared HCR and FISH in control and *C9orf72* patient fibroblasts. Unlike in transiently transfected cells where RNA signal via HCR was predominantly globular and nuclear, endogenous G_4_C_2_ repeats appear primarily as small foci, similar to foci detected by FISH. Previous studies observed upwards of 35% of *C9orf72* patient fibroblasts contained at least 1 G_4_C_2_ repeat foci (Liu et al. 2017), although signal specificity was not established. Here, after normalizing to control fibroblast signal, we observed only half the number of G_4_C_2_ repeat positive cells previously reported in three different expansion cell lines (C9-C1, C9-C2, C9-C3). However, our number of foci/foci positive cell is in agreement with previously published work (Figure 3E) (Liu et al. 2017; DeJesus-Hernandez et al. 2017). Using HCR, we detected >2x more G_4_C_2_ positive cells than with FISH (Figure 3A-B, D). Importantly, we also observed a significant increase in the number of foci/foci positive cell (Figure 3E). Intriguingly, while FISH primarily detected foci in the nucleus of G_4_C_2_ repeat expansion cell lines, HCR was able to detect cytoplasmic foci in ~40% of G_4_C_2_ repeat positive cells (Figure 3F). Previous studies observed cytoplasmic foci only ~10% of the time (Mizielinska et al. 2013). To confirm that the signal we were observing was from RNA, we treated the G_4_C_2_ repeat expansion fibroblasts with DNase or RNase before HCR and found the HCR signal was sensitive to RNase and resistant to DNase (Figure 3C).

**Figure 3.**
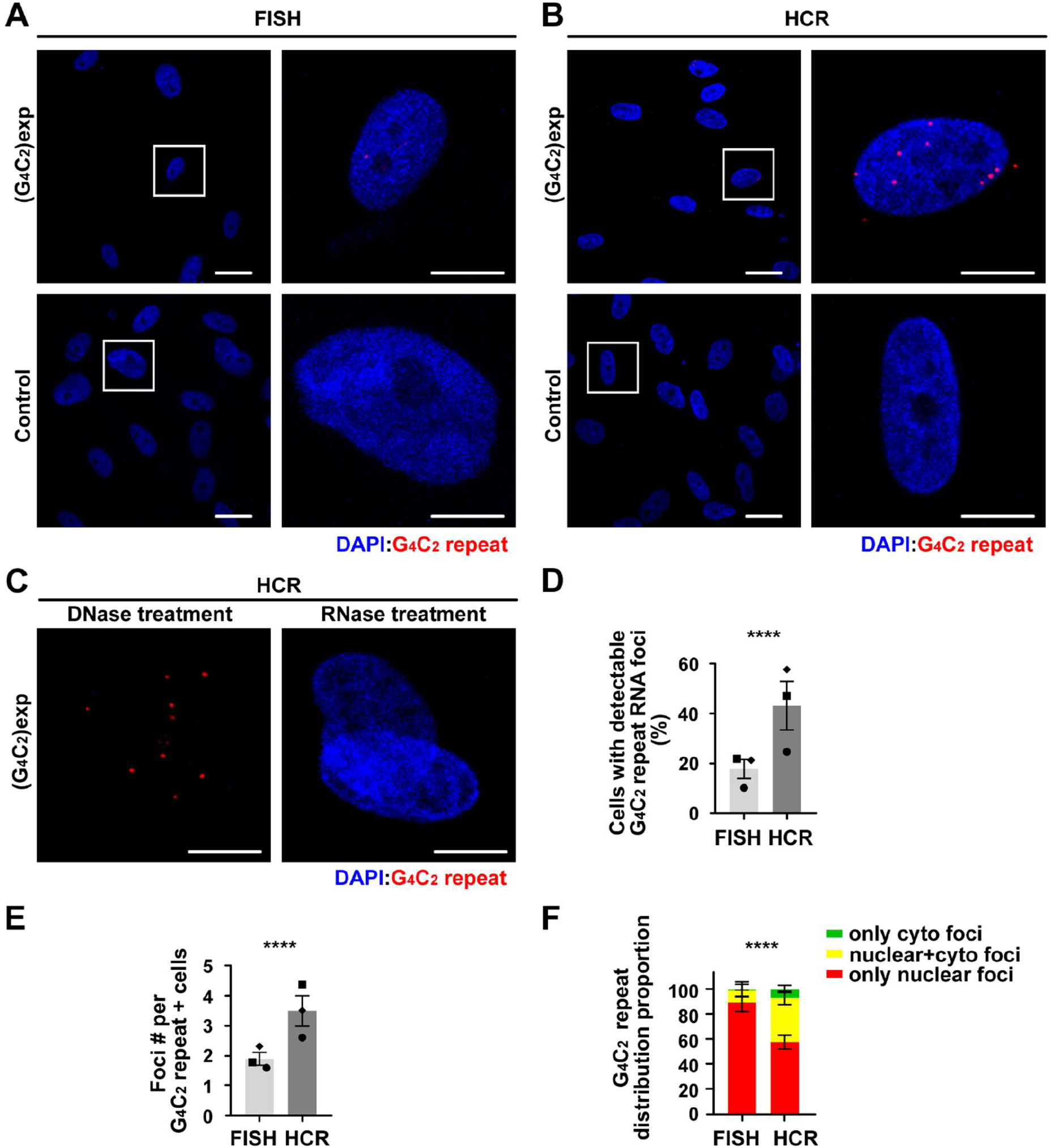
HCR is more sensitive than FISH in detecting endogenous G_4_C_2_ repeat signal from *C9ORF72* ALS-FTD patient fibroblasts. **A-C)** *C9ORF72* ALS-FTD patients and control fibroblasts by FISH and HCR, and treated with DNase and RNase prior to HCR. Higher magnification images of the boxed areas are shown on the right. **D)** Quantification of cells with detectable G_4_C_2_ repeat RNA foci in total cells for fibroblasts. FISH: N=577; HCR: N=690. **E-F)** Quantification of RNA foci number per RNA+ cell and the distribution for RNA foci. The dot shapes represent individual cell lines, which were each assayed in triplicate. FISH: N=106 and HCR: N=305 for E-F). Error bars in D, E indicate SEM, and in F shows 95% CI. Statistics: two-way ANOVA for D and E, Chi-square test for F. \*\*\*\**p*<0.0001. Scale bar=50 μm in left for A and B; 20μm in right for A and B, and C.

We next compared FISH and HCR in control and *C9orf72* patient brain tissue. We looked for foci in cerebellum and frontal cortex, as those regions have been previously shown to have G_4_C_2_ repeat RNA foci and feature evidence of disease pathology (DeJesus-Hernandez et al. 2017; Mizielinska et al. 2013). After normalizing signal to controls, we detected G_4_C_2_ repeat positive cells from three of the C9 brains (C9-B1, C9-B2 and C9-B3) using FISH. However, signal was only detected in about 1% of cells. In contrast, when HCR was performed on these same brain samples, more than 30% of cells were positive for G_4_C_2_ repeat RNA (Figure 4A-B, D). These foci were absent when tissue was RNase treated, and remained when tissue was DNase treated, strongly supporting that these are RNA foci (Figure 4C). Together, these data indicate that HCR can be useful in detection of low abundant endogenous G_4_C_2_ repeat RNA.

**Figure 4.**
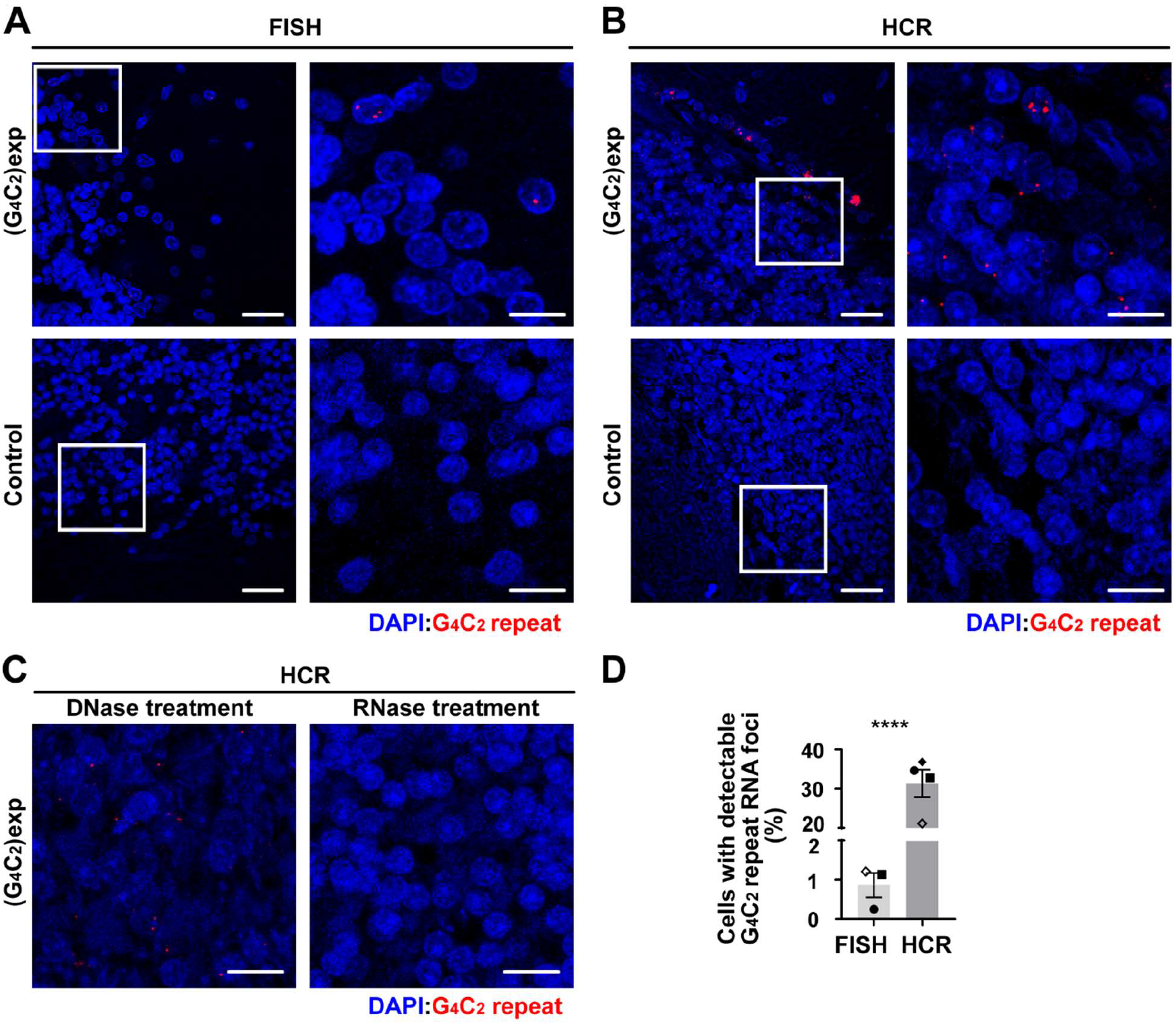
HCR is more sensitive than FISH in detecting endogenous G_4_C_2_ repeat signal from *C9ORF72* ALS-FTD patient brains. **A-C)** FISH **(A)** and HCR **(B)** of *C9ORF72* ALS-FTD patient and control cerebellum, and **(C)** treated with DNase and RNase prior to HCR. Higher magnification images of the boxed areas are shown on the right. **D)** Quantification of cells with detectable G_4_C_2_ repeat RNA foci in total cells for brains ± SEM. The dot shapes indicate different brain samples. The black dots are from cerebellum and unfilled dots are from frontal cortex. FISH: N=2976; HCR: N=3285. Statistics: two-way ANOVA for D. \*\*\*\**p*<0.0001. Scale bar=50μ m in left for A and B; 20 μ m in right for A and B, and C.

We performed a similar experiment in control and FXTAS patient fibroblasts. With the sense CGG repeat HCR probe, we found CGG repeat signal was readily detectable not only in FXTAS patient cells lines, but also in premutation carriers and control lines. The pattern of the detected signal appeared predominantly nucleolar similar to prior reports in FXTAS brain tissue (Tassone, Iwahashi, and Hagerman 2004; Sellier et al. 2013; Sellier et al. 2010). We did not observe any HCR signal in these same cell lines when we used the antisense CCG repeat HCR probe or after RNAse treatment, suggesting this signal was primarily CGG RNA-mediated (Supplemental Figure 2A). These results indicate that the CGG repeat signals are CGG RNA-specific but not *FMR1* CGG repeat expansion-specific. We next evaluated HCR on CGG repeats in frontal cortex and hippocampal sections as these have been previously shown to express the CGG RAN product, FmrpolyG (Krans et al. 2019). Similar to fibroblasts, we observed nucleolar-like staining in both control and FXTAS patient brain tissue with both the CGG FISH and HCR probes. However, the signal was stronger and occurred in a higher percentage of neurons in FXTAS samples (Supplemental Figure 2B).

As the control cell lines still contain ~20-30 CGG repeats, we reasoned that the probe could still be specifically binding to *FMR1* RNA. To investigate this further, we performed HCR in a transcriptionally silenced FXS iPSC line with 800 repeats, a control iPSC line with ~30 CGG repeats and then compared their signal intensity to an unmethylated full mutation line (TC-43) with a large (270) transcriptionally active CGG repeat expansion that supports RAN translation (Haenfler et al. 2018; Rodriguez et al. 2020). We observed significant staining with the CGG HCR probe in all three cell lines that was both diffuse in the cytoplasm, as well as localized to the nucleolus (Supplemental Figure 2C) However, similar to observations in FXTAS brain, we saw enhanced signal in the unmethylated full mutation line compared to the WT and methylated FXS line, suggesting that this enhanced signal was due to CGG repeat expansions within *FMR1* (Supplemental Figure 2C-E).

### G4C2 repeats accumulate in the nucleus in response to cellular stress

The ability to readily detect endogenous nuclear and cytoplasmic G_4_C_2_ repeat RNA foci using HCR is potentially useful for exploring its roles in disease pathogenesis. Our lab and others have observed that expression of G_4_C_2_ repeat containing reporters induces stress granule (SG) formation and the integrated stress response (ISR), and considerable evidence now suggests that this process can contribute to neurodegeneration (Green et al. 2017; Sonobe et al. 2018; Zhang et al. 2018; Li et al. 2013). Moreover, exogenous ISR activation through a variety of methods triggers a selective enhancement of RAN translation from both CGG and G_4_C_2_ repeats in transfected cells and neurons (Green et al. 2017; Westergard et al. 2019; Sonobe et al. 2018; Cheng et al. 2018). To investigate the behavior of endogenous G_4_C_2_ repeat RNA and foci in response to stress, we treated *C9orf72* patient fibroblasts with sodium arsenite (SA) or vehicle. SA treatment for one or two hours led to significant increase in the total number of cells with visible G_4_C_2_ repeat foci. Moreover, there was a marked re-distribution of these foci into the nucleus and out of the cytoplasm (Figure 5A-C). This same SA-induced nuclear re-distribution of G_4_C_2_ repeat foci was also observed in *C9orf72* patient derived neurons (Figure 6A-C).

**Figure 5.**
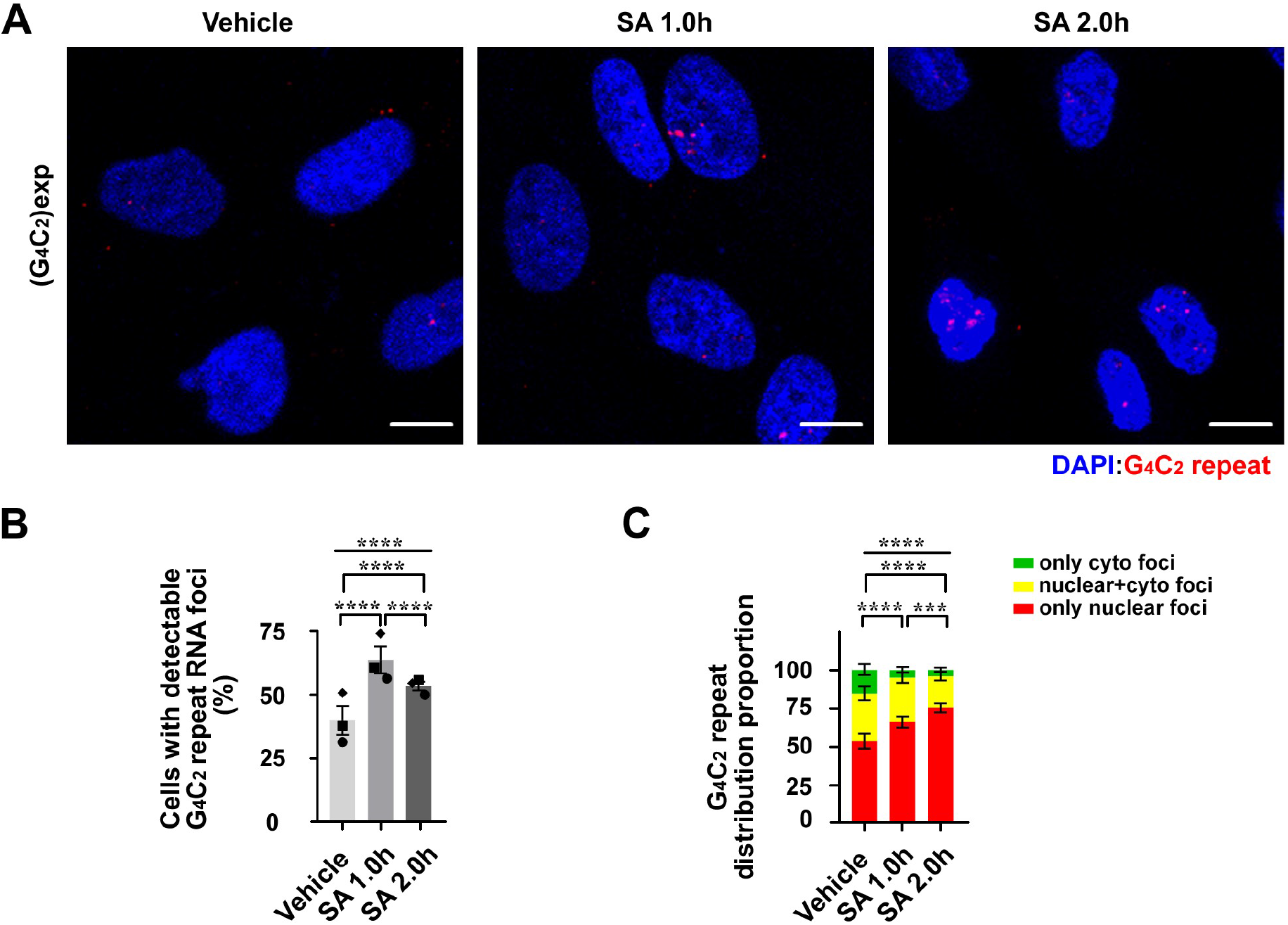
G_4_C_2_ repeats redistribute into the nucleus during stress in *C9ORF72* ALS-FTD patient fibroblasts. **A)** *C9ORF72* ALS-FTD patient fibroblasts treated with H_2_O (vehicle) or sodium arsenite (SA) for the indicated times prior to fixation and HCR. **B)** Quantification of cells with detectable G_4_C_2_ repeat RNA foci in total cells ±SEM. The dot shapes represent different G_4_C_2_ repeat expansion cell lines. N=1130 for vehicle, 1120 for SA 1.0h, and 1482 for SA 2.0h. **C)** Distribution of G_4_C_2_ repeat RNA foci with or without SA treatment ± 95% CI. N=400 for vehicle, 679 for SA 1.0h, and 775 for SA 2.0h. Statistics: two-way ANOVA for B, Chi-square test for C. \*\*\**p*<0.001, \*\*\*\**p*<0.0001. Scale bar=10 μ m in A.

**Figure 6.**
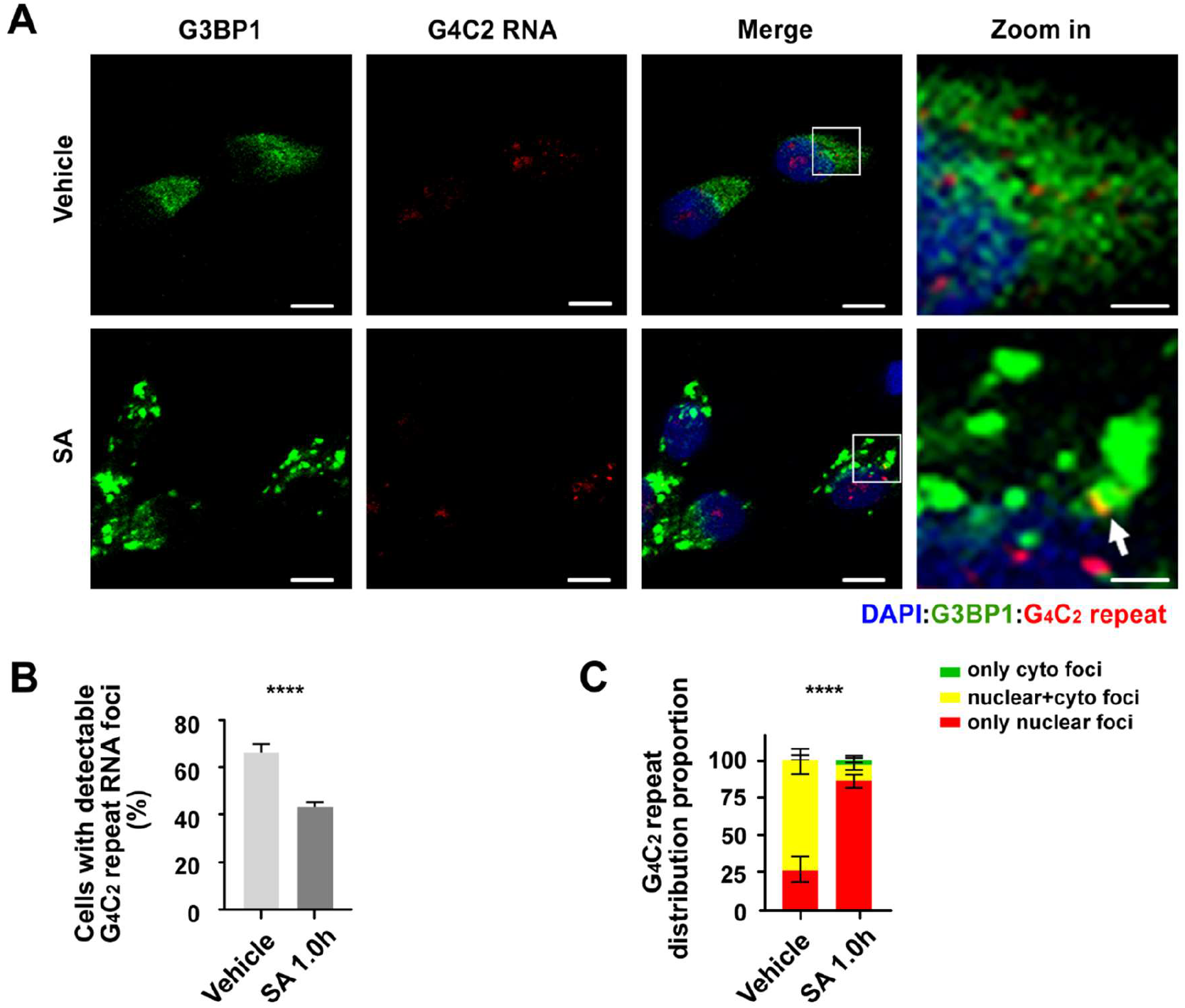
G_4_C_2_ repeats redistribute in the nucleus during stress in *C9ORF72* ALS-FTD patient neurons. **A)** *C9orf72* patient derived neurons treated with H_2_O or SA prior to HCR-ICC. G3BP1= stress granule marker. A higher magnification of the boxed areas are shown on the right. White arrows indicate rare colocalization events between cytoplasmic G4C2 repeats and G3BP1. **B)** Cell with detectable G_4_C_2_ repeat RNA among total neurons treated with or without SA ±SEM. N=156 for vehicle and 535 for SA 1.0h. **C)** Quantification of the distribution of repeat RNA in all repeat positive neurons ± 95% CI. N=104 for vehicle and 233 for SA 1.0h. Statistics: unpaired t test for B and Chi-square test for C. \*\*\*\**p*<0.0001. Scale bar=10 μ m in A except for the zoom in images (2.5 μ m).

RNAs typically move into SGs and become translationally silenced in response to SA stress (Kedersha et al. 2005). In contrast, G_4_C_2_ repeat RNAs remain translationally competent after stress induction. We therefore assayed whether G_4_C_2_ repeat RNA foci localized to SGs in response to stress. Consistent with their retained translational competency, we did not observe significant co-localization of G_4_C_2_ repeat foci with the SG marker, G3BP1 (Figure 7A-B), although rare co-localization events were observed.

**Figure 7.**
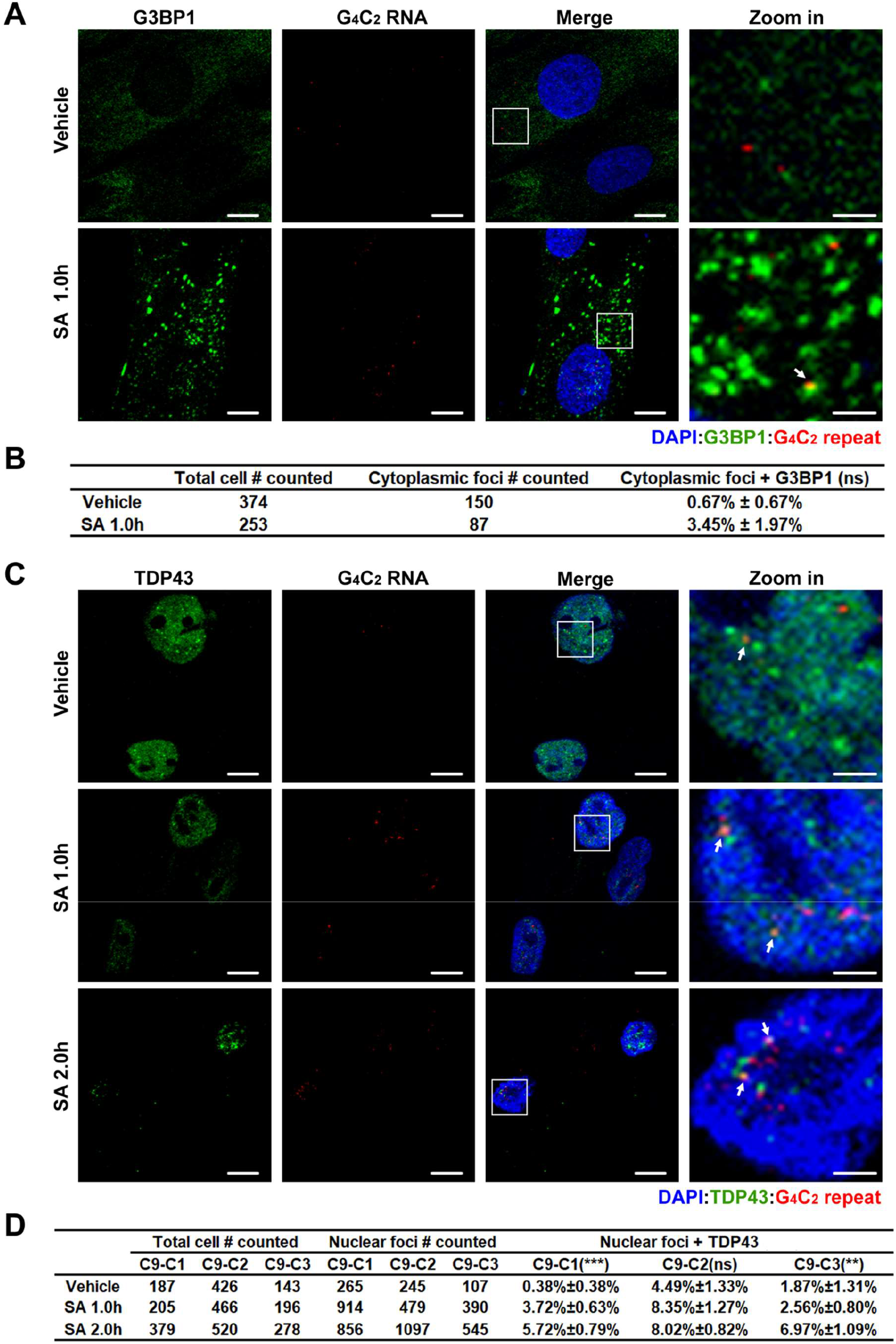
G_4_C_2_ repeats rarely co-localize with cytoplasmic G3BP1 or nuclear TDP43 during stress. **A)** *C9orf72* patient derived fibroblasts treated 1 hour with H_2_O or SA prior to HCR-ICC. G3BP1= stress granule marker. Higher magnification images of the boxed areas are shown on the right. White arrows indicate co-localization of G4C2 repeat and G3BP1. **B)** Quantification of cytoplasmic G_4_C_2_ repeat foci co-localization with G3BP1 as a fraction of all cytoplasmic G_4_C_2_ repeats. **C)** *C9orf72* patient derived fibroblasts treated with H_2_O or SA (1 hour, 2 hours) prior to HCR-ICC. TDP43= nuclear body marker. White arrows indicate co-localization of G_4_C_2_ repeat and TDP43. **D)** Quantification of nuclear G_4_C_2_ repeat foci colocalization with TDP43 as a fraction of all observed nuclear G_4_C_2_ repeat foci. % shown as mean±SEM. Statistics: unpaired t test for B and one-way ANOVA for D. \*\**p*<0.01, \*\*\**p*<0.001. ns: not significant. Scale bar=10 μ m in A and C except for zoomed images (2.5 μ m).

Integrated Stress response activation induces the formation of nuclear stress bodies. These TDP-43 positive structures are thought to contribute to ALS disease pathogenesis (Wang et al. 2020). Given the greater nuclear distribution of G_4_C_2_ repeat RNA after stress induction, we evaluated whether there was any significant overlap between endogenous G_4_C_2_ repeat RNA and TDP43, a critical factor in C9orf72 pathology and ALS pathogenesis as well as a robust marker for nuclear bodies (Duan et al. 2019; Wang et al. 2020). Our untreated C9 fibroblasts appeared to have small TDP-43 nuclear bodies, indicative of them being inherently stressed. Upon 2 hours of SA induction we saw a decrease in diffuse nuclear TDP43, and an increase in TDP-43 nuclear foci size. Overall, nuclear TDP-43 showed limited (0.38-4.49%) co-localization with G_4_C_2_ repeat RNA foci at baseline. After SA induction, there was a significant increase in this co-localization, but it remained modest (2.56%~8.35%) (Figure 7C-D). Taken together, these studies suggest that nuclear retention or re-distribution of G_4_C_2_ repeat RNA foci in C9orf72 fibroblasts in response to stress is not predominantly driven by either SG or nuclear body association and may instead reflect nucleocytoplasmic transport defects elicited by stress pathway activation (Mahboubi et al. 2013; Zhang et al. 2018; Burgess et al. 2011; Patterson, Wood, and Schisa 2011).

## Discussion

Repeat RNA and the formation of RNA-protein and RNA-RNA condensates are thought to act as significant factors in the pathogenesis of multiple repeat expansion disorders. However, traditional detection techniques such as FISH are often limited in their sensitivity which may cloud the roles of such repeat RNAs in disease-relevant processes. Here we used a highly sensitive RNA in situ amplification method, HCR, to readily detect low expressing endogenous GC-rich repeat expansions in both patient cells and tissues. This non-proprietary method provided significantly enhanced sensitivity over RNA FISH probes with retained specificity. This increased sensitivity allowed for greater detection and appreciation of nuclear and cytoplasmic foci in patient cells. Moreover, we demonstrated that G4C2 repeat RNA foci accumulate in the nucleus in both patient fibroblasts and neurons in response to cellular stress. This tool should prove useful to the field in explorations of endogenous repeat RNA behaviors and pathology in both repeat expansion disorders and model systems.

Previously established techniques for detecting RNA in situ have limitations. FISH is limited by RNA copy number, and probe specificity, while the use of MS2 and PP7 binding sites to detect low abundant RNAs is only applicable to exogenous gene expression, or in cases where these tags were inserted via CRISPR (Tantale et al. 2016; Spille et al. 2019; Ferguson and Larson 2013; Young, Jackson, and Wyeth 2020). Recently, an alternative RNA in situ amplification method, Basescope™, was shown to improve detection of endogenous G_4_C_2_ repeats in patient tissue (Mehta et al. 2020). HCR and Basescope™ are comparable on a number of fronts. Namely, they both can be combined with IHC, have extensive signal amplification capacity, and in the case of newer HCR versions, utilize split probes to eliminate nonspecific signal (Choi et al. 2018). However, HCR lends itself as a more universally applicable approach for a number of reasons, including minimal optimization needed, fewer steps involved, flexibility in probe stringency, and the use of fluorescent probes to allow for combined HCR-IF with multiple probes and/or antibodies. Furthermore, unlike Basescope™, HCR v2 reagents are not proprietary, allowing for in-house production and lower cost. However, while the flexibility of fluorescent probe choice makes HCR more adaptable in a variety of experimental settings, the chromogenic properties of Basescope™ may allow for better coupling with tissue stains for pathology purposes. Thus, both serve as valuable tools for detecting and investigating endogenous GC-rich repeats.

While both HCR and Basescope™ are sensitive tools for detecting G_4_C_2_ repeat RNA, we do caution the use of these techniques for detecting CGG repeat RNA. Given the extensive use of CGG RNA FISH probes in the literature, we were surprised to find such high background signal, specifically in control and FXS human samples. The staining pattern with the CGG probe was also vastly different from the foci typically observed for GC-rich repeat expansions. In fibroblasts, brain samples, and HEK293 cells we observed intense, large nuclear body staining, while iPSCs had weaker nuclear body staining and strong, diffuse cytoplasmic staining. This pattern is largely consistent with prior work using FISH to assay CGG repeat RNA in FXTAS (Sellier et al. 2013; Sellier et al. 2010). That lack of punctate RNA foci makes it difficult to determine what signal is specific to the CGG repeat expansion on *FMR1*. There are 921 human genes which contain ≥6 CGG repeats, and this likely accounts for the high background in human cells (Haenfler et al. 2018; Kozlowski, de Mezer, and Krzyzosiak 2010). Future studies using probes to more specific regions of *FMR1* are needed.

Our HCR G_4_C_2_ probe exhibited significantly better sensitivity and specificity in human cells and tissues. The increased sensitivity of HCR over FISH with this probe allowed us to consistently visualize G_4_C_2_ repeat RNA in the cytoplasm, allowing us to ask questions regarding G_4_C_2_ repeat RNA activities in different subcellular compartments. As a proof of concept, we analyzed G_4_C_2_ repeat cellular localization during stress. We observed no significant co-localization with SGs, but instead, a redistribution of G_4_C_2_ foci into the nucleus. This redistribution could either be caused by an increase in nuclear import, a decrease in nuclear export, or a retention of repeat RNA within sub-nuclear compartments. Nucleocytoplasmic transport is inhibited basally in many neurodegenerative conditions, including C9orf72 ALS/FTD (Rossi et al. 2015). Nucleocytoplasmic transport proteins, including importin-alpha, RanGap and nucleoporins are also recruited into SGs and co-localize with TDP-43 in ALS/FTD mutant cytoplasmic aggregates (Chou et al. 2018; Gasset-Rosa et al. 2019). SG assembly itself inhibits nucleocytoplasmic transport by sequestering factors required for nuclear export, and thus the increased abundance of G_4_C_2_ repeat RNA may be indicative of global nuclear mRNA retention (Mahboubi et al. 2013; Zhang et al. 2018; Burgess et al. 2011; Patterson, Wood, and Schisa 2011). G_4_C_2_ repeat RNA itself is also implicated in nuclear import perturbations via binding to RanGap1 (Zhang et al. 2015), suggesting that its nuclear retention could be not only a cause of but also a contributor to stress-dependent pathology.

Alternatively, repeat RNAs may interact with nuclear stress bodies. These complexes result from nuclear re-localization of heat shock factors, including HSF1 and HSP70, as well as RNA factors, including TDP-43, with satellite III repeat RNAs (Udan-Johns et al. 2014; Metz et al. 2004; Frottin et al. 2019; Yu et al. 2020). We observed a significant increase in co-localization between nuclear TDP-43 and G_4_C_2_ RNA. However, the overall overlap between these two molecules remained modest and of unclear biological significance.

In sum, we describe the application of HCR to the detection of endogenous GC rich repeat RNA. This non-proprietary tool is sensitive, specific and useful in studying endogenous repeat RNA foci dynamics and should prove useful for investigators interested in the behavior of these disease-associated RNA species.

## Methods

### 1) Cell lines culture and Clinical Specimens

MEFs were received from Randal Kaufman (Sanford Burnham Prebys Medical Discovery Institute).and cultured in RPMI1640 with 10% fetal bovine serum (FBS) and 1% penicillin/streptomycin (P/S). FXTAS skin fibroblasts were from Paul Hagerman (UC Davis) and University of Michigan donors. *C9orf72* ALS/FTD skin fibroblasts are from Eva Feldman (University of Michigan). Human derived skin fibroblasts were cultured in high glucose DMEM with 10% FBS, 1% non-essential amino acid (NEAA) and 1% P/S. All cells were cultured at 37°C. Control and FXTAS human brain paraffin sections were obtained from the University of Michigan Brain Bank and are previously described (Todd et al. 2013; Krans et al. 2019). Human derived neurons were generated and differentiated from Elizabeth Tank. Human derived iPSCs were generated and differentiated from Geena Skariah. Details on these specimens were presented in Supplemental Table 1.

### 2) Reporters

The RAN translation reporters 2(CGG)_n_-NL-3xF, (G_4_C_2_)_n_-NL-3xFlag, and (CCG)_60_-NL-3xF were previously published (Kearse et al. 2016; Green et al. 2017; Krans et al. 2019). The (G_2_C_4_)_47_-NL-3xFlag reporter was made by cloning the reporter sequence between NheI and PspOMI in pcDNA3.1+ (Supplemental Table 3). The FMRpolyG_100_-RAN translation reporters was made by cloning in the reporter sequence into pcDNA3.1+ between BamHI and PspOMI (Supplemental Table 3).

### 3) Transfection

MEFs were transfected according to manufacturer’s protocol (114-15, Polyplus transfection). In brief, MEFs were suspended and incubated with3:1 Jetprime to 0.25 μg plasmid DNA at 37°C for 20 minutes, then seeded to chamber slides. Cells were fixed 24 hours later for ICC-HCR.

### 4) Stress treatment

Fibroblasts and neurons were treated with 500nM sodium arsenite (SA) or vehicle (H_2_O) at indicated time points then washed 1x in PBS and fixed for ICC-HCR.

### 5) ICC

The ICC was performed after fixing cells and prior to performing HCR, using a modified protocol to one previously described. In brief, following a fix in 4% PFA, cells were further fixed O/N in 70% ethanol, and rehydrated with 1xPBS for 1 hour, prior to addition of antibodies. Cells were then fixed again in 4% PFA for 10 minutes following the final washes after the secondary to fix the secondary antibodies in place. The following antibodies were used: Flag M2 (1:100, F1804, Sigma), GFP (1:500, ab6556, abcam), G3BP1(1:200, BDB611127, BD Bioscience/Fisher), TDP43(1:100, 10782-2-AP, Protein Tech), Alexa Fluor 488 goat anti-mouse IgG (1:500, A11029, Invitrogen) and anti-rabbit IgG (1:500, A11008, Invitrogen).

### 6) FISH

Fluorophore TYE665 labeled locked nuclear acid (LNA) (C_4_G_2_)_6_ and (CCG)_8_ probes (Supplemental table 2) were synthesized by Qiagen. The FISH protocol was adapted from (DeJesus-Hernandez et al. 2017) and used HCR buffers (Choi, Beck, and Pierce 2014b; Choi et al. 2011). In brief, MEFs and fibroblasts, were washed with 1x PBS-MC, fixed in 4% PFA for 10 min, washed 3 times with 1x PBS and permeabilized in 0.1% TritonX-100 for 15 min. Cells were washed with 1x PBS, then dehydrated with 70% ethanol for 1min, 95% for 2min, then 100% twice for 2min. Cells were air-dried, then preheated at corresponding hybridization temperature (see below) in probe hybridization buffer (50% formamide, 5x sodium chloride sodium citrate (SSC), 9 mM citric acid (pH 6.0), 0.1% Tween 20, 50μg/mL heparin, 1x Denhardt’s solution, 10% dextran sulfate) for 30 min. The FISH probes were denatured at 80°C for 2 min before immediately snap cooling in cold hybridization buffer to prevent intermolecular annealing. Cells were incubated in hybridization buffer with 8nM, 32nM or 64nM probes at 71°C for (C_4_G_2_)_6_ and 66°C for (CCG)_8_ for 12-16hrs, washed 4 times for 5min in 5xSSCT (5x SSC, 0.1% Tween 20) at room temperature (RT), then mounted with ProLong Gold antifade mountant with DAPI.

For human brain paraffin sections, slides were first deparaffinized with xylenes and then rehydrated from 100% ethanol, 95% ethanol, 70% ethanol, 50% ethanol to DEPC treated H_2_O. Rehydrated slides were incubated in 0.3% Sudan black for 5 min followed by proteinase K treatment at RT for 10 min. Permeabilization, preheating, hybridization with probes, washing and mounting are the same as above.

### 6) HCR

The initiator probes (CCCCGG)_6_ and (CCG)_10_, were synthesized by OligoIDT. The fluorophore 647 labeled hairpin probes (B1H1 and B1H2) (Choi, Beck, and Pierce 2014b) were synthesized by Molecular Instruments. For transfected MEFs and fibroblasts, cells processed according to Molecular Instrument’s protocol. In brief, cells were fixed in 70% cold ethanol overnight at 4°C. When HCR was performed following ICC, this step occurred prior to ICC.,Cells were then preheat in hybridization buffer at 45°C for 30min, and incubated with 0.8nM, 4nM or 8nM initiator probe ((CCCCGG)_6_ or (CCG)_10_) at 45°C incubator overnight. Cells were washed 4 times for 5 minutes each with pre-warmed probe wash buffer (50% formamide, 5x SSC, 9 mM citric acid (pH 6.0), 0.1% Tween 20, 50μg/mL heparin) at 45°C and 2 time for 5min with 5x SSCT at RT. Cells were incubated with snap cooled hairpins B1H1 and B1H2 at room temperature for 12-16 hours. The concentrations of each hairpin (0.375 pmol, 1.875 pmol, and 3.75 pmol per well in an 8 well chamber slide) was proportional to the amount of initiator probe used (0.8nM, 4nM or 8nM). Cells were washed 5 times for 5 minutes at RT with 5x SSCT, then mounted with ProLong Gold antifade mountant with DAPI.

For human derived brain section, samples were processed according to Molecular Instrument’s protocol. In brief, Histo-Clear II was used to deparaffinize tissue, then samples were rehydrated and treated with proteinase K as described for FISH., Slides were washed in 1x TBST, incubated in 0.2N HCL for 20min at RT, washed5 times in 5xSSCT, then incubated in 0.1M triethanolamine-HCR (pH 8.0) with acetic anhydride for 10 min, and washed in 5x SSCT for 5min. Slides were preheated and incubated in hybridization buffer with probes at 45°C in humidity chambers for 12-16 hrs. The remaining steps weree the same as above for HCR in cell culture.

### 8) Imaging and analysis

Images were taken on an Olympus FV1000 confocal microscope equipped with a 40x oil objective (60x for iPSC images). ImageJ software was used to analyzed the exported images. The backgrounds for stained proteins and repeat RNA signals were normalized to non-transfected group or controls unless indicated. Total cell number, protein stained cell number, RNA positive cell number, and foci number and distribution per cell were all manually counted. Signal intensities for iPSC images (Supplemental Figure 2E) were calculated as mean intensity/area using ImageJ. For transfected MEFs, RNA intensity was graded into high, medium and low signal intensity. The ratio of repeat positive cells was calculated as number of RNA positive cells to total cells. The ratio of foci number per cell was calculated as foci number in all repeat positive cells divided by all repeat positive cells. The repeat distribution was expressed as proportion of cells with foci (only nuclear, only cytoplasmic and both) among all repeat positive cells. The relationship between repeat foci and SGs was analyzed as the proportion of co-localization of cytoplasmic G_4_C_2_ repeat foci with G3BP1 granule to total cytoplasmic G_4_C_2_ repeat foci. Similarly, the relationship between repeat foci and NBs was analyzed as the proportion of co-localization of nuclear G_4_C_2_ repeat foci with TDP43 granule to all nuclear G_4_C_2_ repeat foci.

### 9) Statistical analysis

All statistical analyses were performed in GraphPad Prism software. Chi-square test was applied for categorical data, including amount of stained protein and repeat cells in transfected MEFs, repeat intensity in transfected MEFs, and distribution of repeat foci in cells. Unpaired t-test, one-way ANOVA and two-way ANOVA were performed to analyze continuous data, including number of detectable repeat foci, foci number per cell, and the co-association rates between repeat RNA and SG or NB markers. We designated *P*<0.05 as our threshold for significance.

---

**Supplemental Table 1.** Information for human derived samples.

| Sample ID | Lab ID | Type | Diagnosis | CGG/G<sub>4</sub>C<sub>2</sub> repeat NO. |
| --- | --- | --- | --- | --- |
| C9-C1 | 499 | fibroblasts | C9-ALS/FTD | G<sub>4</sub>C<sub>2</sub>=1180 |
| C9-C2 | 733 | fibroblasts | C9-ALS/FTD | G<sub>4</sub>C<sub>2</sub>=1100 |
| C9-C3 | 953 | fibroblasts | C9-ALS/FTD | G<sub>4</sub>C<sub>2</sub>=1230 |
| Control-C1 | C1 | fibroblasts | Control | CGG=31 |
| Control-C4 | C4 | fibroblasts | Control | CGG=22 |
| C9-B1 | exp198 | Brain-cb | C9-ALS/FTD | Expanded (RP-PCR) |
| C9-B2 | exp1233 | Brain-cb | C9-ALS/FTD | Expanded (RP-PCR) |
| C9-B3 | exp1350 | Brain-cb<br>Brain-cortex | C9-ALS/FTD | Expanded (RP-PCR) |
| Control-B1 | cont1033 | Brain-cb<br>Brain-cortex | Control | n.d. |
| Control-B2 | cont1533 | Brain-cb | Control | n.d. |
| FXTAS-1 | F1 | fibroblast | FXTAS | CGG=122 |
| FXTAS-3 | F3 | fibroblast | FXTAS | CGG=97 |
| Premutation-3 | P3 | fibroblast | Premutation carrier | CGG=81 |
| Premutation-4 | P4 | fibroblast | Premutation carrier | CGG=70 |
| FXTAS-B1 | 1906 | Brain-hippo | FXTAS | CGG=102 |
| FXTAS-B2 | T345 | Brain-hippo<br>Brain-cortex | FXTAS | CGG=90 |
| Control-B3 | 340 | Brains-cortex | Control | n.d. |
| Control-B4 | T357 | Brains-cortex | Control | n.d. |
| C9-neuron | 882 | Neuron | C9-ALS/FTD | G<sub>4</sub>C<sub>2</sub>>44 |
| Control-neuron | 1021 | Neuron | Control | G<sub>4</sub>C<sub>2</sub>=14 |
| WT | wt4-6 | iPSC | Control | CGG=30 |
| TC43 | TC43 | iPSC | unmethylated FM | CGG=270 |
| FXS | 139-2 | iPSC | FXS | CGG=~800 |

**Supplemental table 2.**
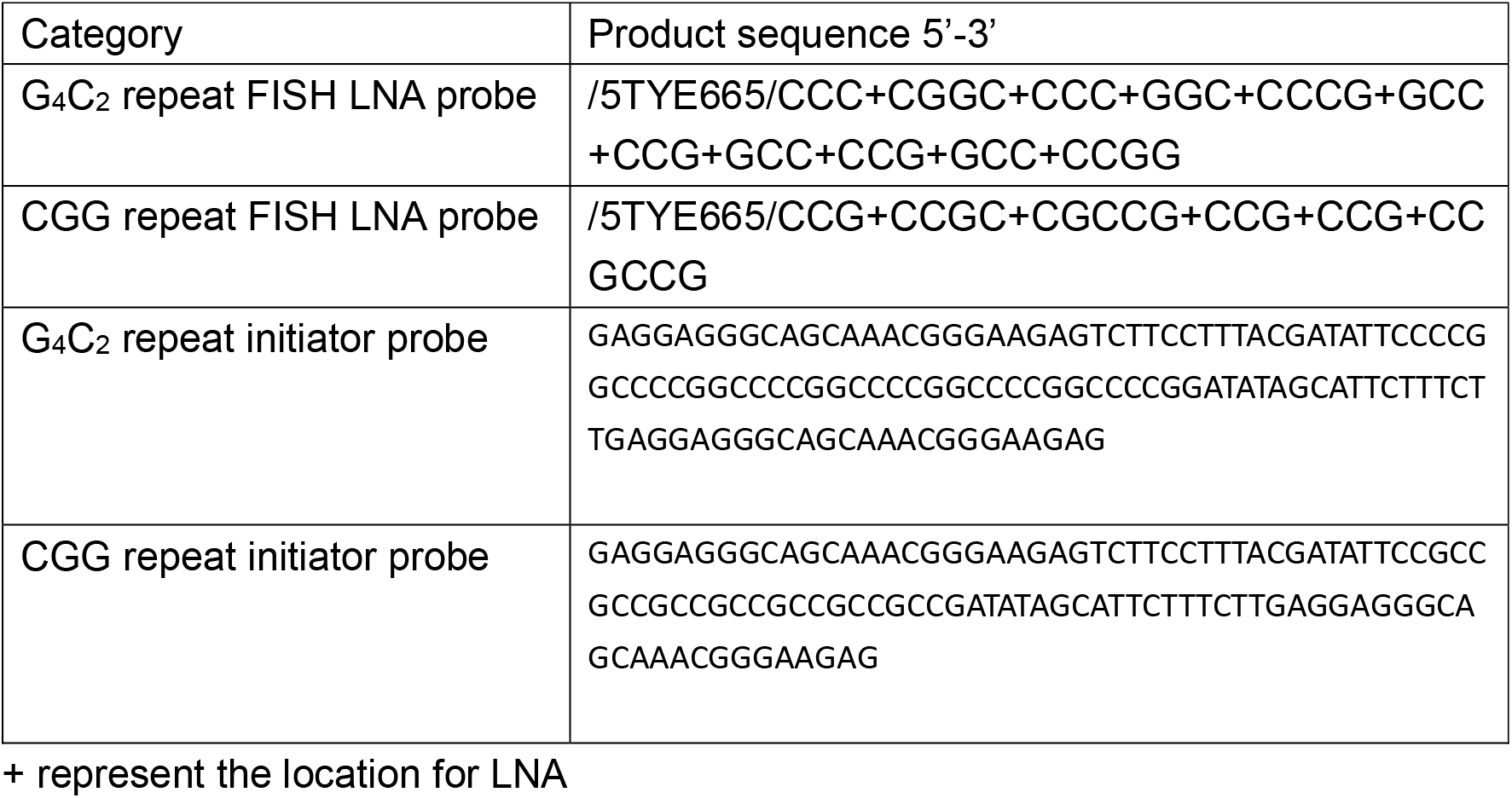
LNA FISH probe and HCR probe sequences

**Supplemental table 3.**
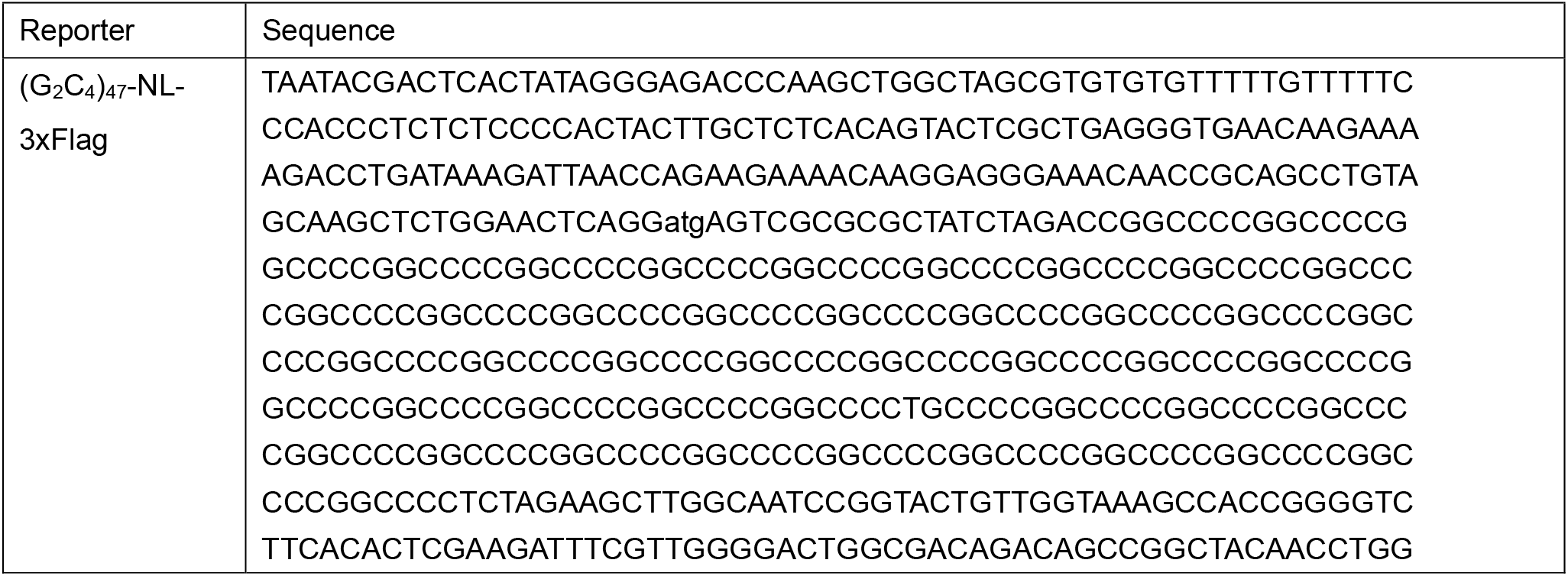
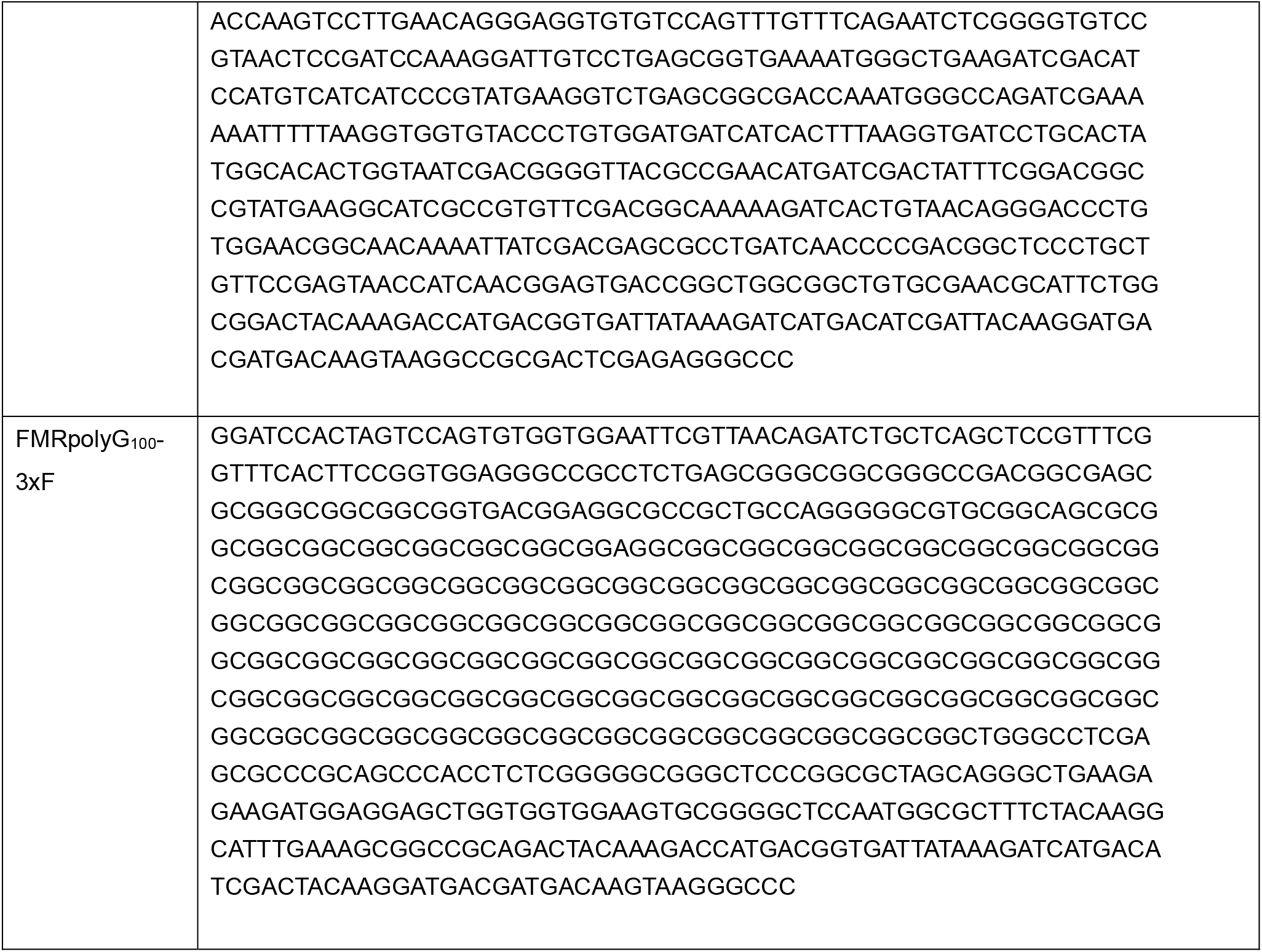
Reporter Sequences

## Acknowledgements

The authors thank Geena Skariah for providing UFM, FXS, and WT iSPC cells for analysis, Amy Krans for assistance with C9orf72 ALS/FTD and FXTAS brain tissues, and Michael Sutton for the use of his confocal microscope Special thanks to David Turner, and Harry Choi for their inspiration and guidance in designing HCR probes. Brain samples were obtained from the University of Michigan Brain Bank with help from Steven Goutman and Eva Feldman in the ALS Center at the University of Michigan who also assisted with provision of fibroblasts.

## Funding

This work was funded by grants from the NIH (P50HD104463, R01NS099280 and R01NS086810 to PKT and T32NS00722237 to MRG) and the VA (BLRD BX004842 to PKT). MRG was supported by the A2A3 Susan Ross Award. YZ was supported by the Chinese Scholarship Council and by private philanthropic support to PKT.

---

**Supplemental Figure 1.**
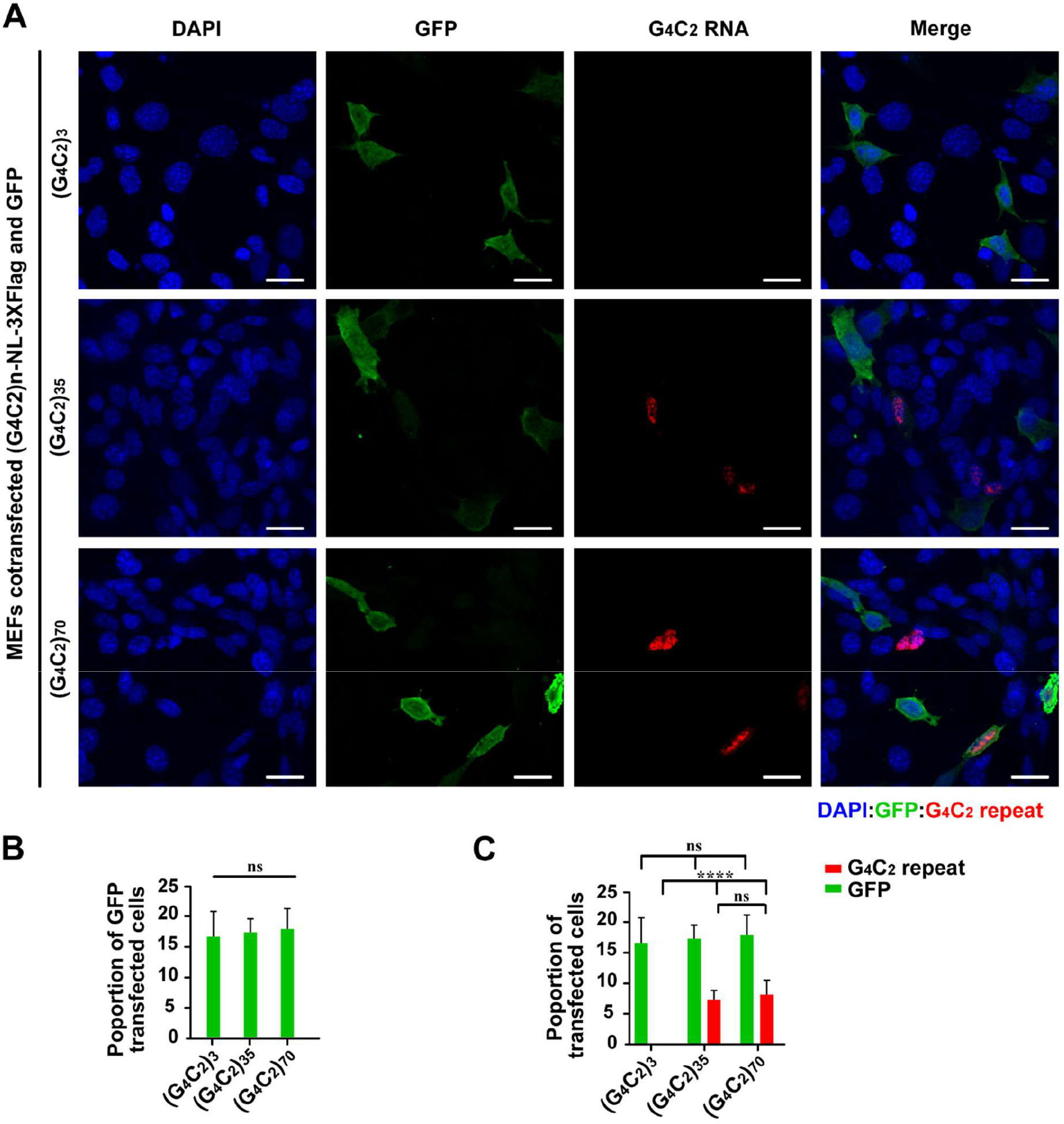
Transfection efficiencies for MEFs co-transfected with GFP and G_4_C_2_ repeats. **A)** MEFs co-transfected with GFP and G_4_C_2_ repeat constructs with increasing repeat sizes. **B-C)** Quantification of cells with detectable GFP and G_4_C_2_ repeat RNA foci signal in total cells. (G_4_C_2_)_3_: N=313; (G_4_C_2_)_35_ N=575; (G_4_C_2_)_70_ N=514. Statistics: one-way ANOVA for B, one-way ANOVA and unpaired t test for C. \*\*\*\**p*<0.0001, ns: not significant. Scale bar=50 μ m in A.

**Supplemental Figure 2.**
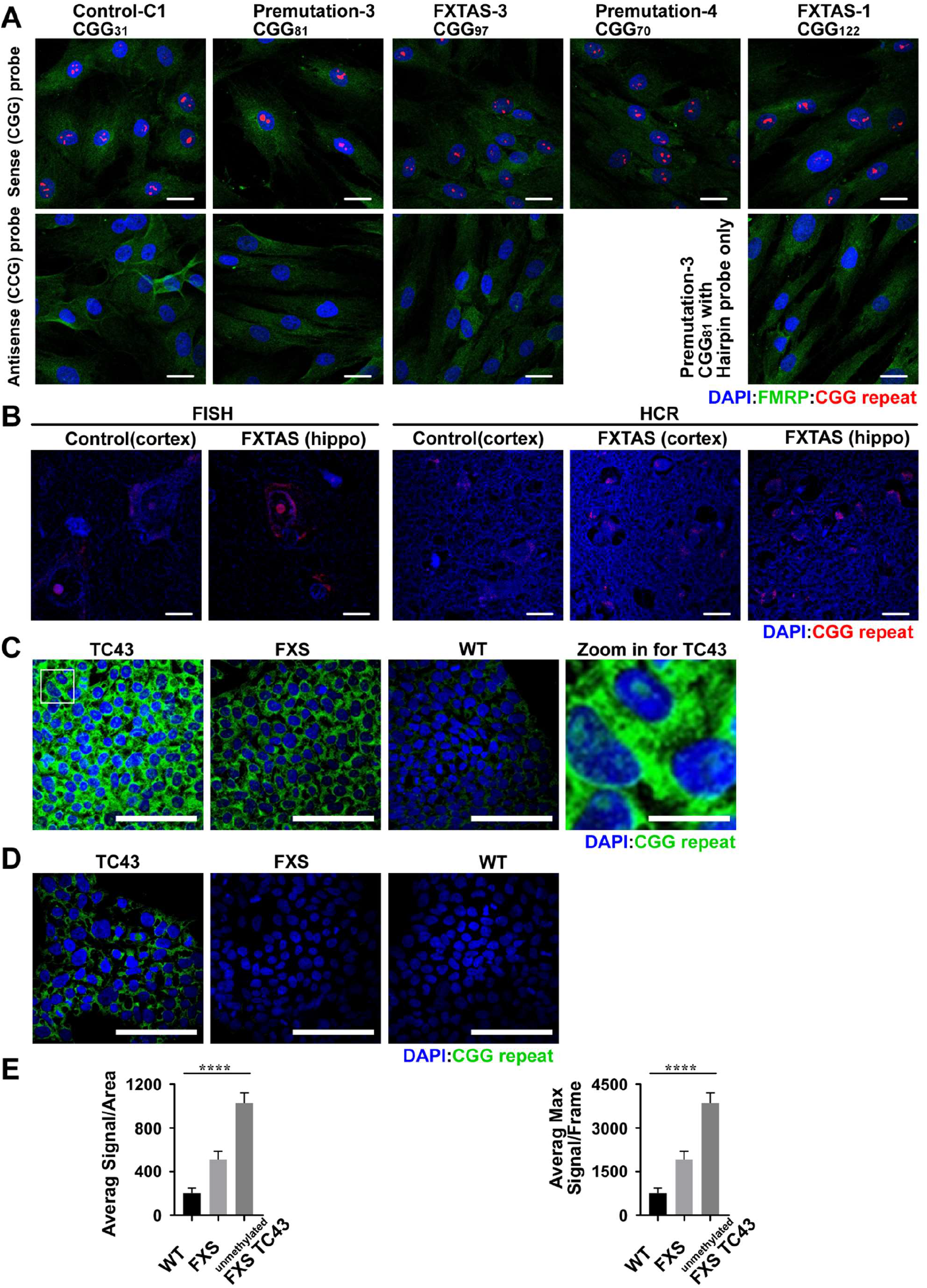
Endogenous CGG repeat is highly expressed in human cells. **A)** HCR of FXTAS patient, permutation carrier and control fibroblasts. **B)** FISH and HCR of FXTAS patient and control brains. **C-D)** iPSCs/hESCs derived from an unmethylated CGG expansion carrier (TC-43; 270 CGG repeats, transcriptionally active), fully methylated embryonic cells (FXS, 800 CGG repeats, transcriptionally silent) and a control patient (30 CGG repeats) were assayed by CGG HCR. Panel C is without background subtraction, demonstrating signal in all cells. Panel D is after subtraction of signal detected in FXS line, revealing significant signal above background only in the actively transcribed expanded CGG repeat line. **E)** Quantification of CGG repeat RNA foci signal in each cell line of figure **(C)** *left:* average signal/frame, *right:* maximum signal/frame. Statistics of one-way ANOVA for E. n=15, \*\*\*\**p*<0.0001. Scale bar=50 μ m in A; 20 μ m in B; 100 μ m in C and D except for the zoom in picture of C (20 μ m).

## Notes

### Competing Interest Statement

The authors have declared no competing interest.

